# Global picoplankton biogeography revealed by metagenomic and climatic data integration

**DOI:** 10.1101/2024.11.23.624595

**Authors:** Vinícius W. Salazar, Heroen Verbruggen, Vanessa Rossetto Marcelino, Kim-Anh Lê Cao

## Abstract

Microbial plankton play fundamental roles in biogeochemical cycles, driving nutrient cycling that influences the global climate and supports life on Earth. Picoplankton are the smallest and most abundant planktonic organisms. The distribution and ecology of these organisms is determined by environmental factors and their biogeography is largely shaped by basin-scale patterns of physicochemical composition of ocean waters. The increased availability of high-throughput sequencing data of microbial communities has enabled the description of how the global oceans are partitioned into distinct microbial biogeographical provinces. However, the key attributes associated with such provinces are still unclear. Here we present a model of picoplankton biogeography based on 1454 metagenomes from multiple sampling consortia, resulting in the largest integrated surface ocean metagenome analysis to date. We identify ten distinct groups based on metagenomic dissimilarity, divided into three categories: polar (Arctic and Antarctic), temperate (coastal temperate, temperate/subtropical transition, oceanic temperate, Mediterranean-like) and tropical (tropical low nutrient, tropical high nutrient, subtropical oceanic gyres). Using machine learning and omics data integration techniques, we predict province areas across the surface oceans and describe their environmental, taxonomical, and functional features. We quantify the relationship between environmental factors and each biogeographical province, identify their main representative taxa and the importance of carbon degradation and antimicrobial resistance pathways in functional community composition, and discuss implications for establishing a model for global picoplankton biogeography.

## Introduction

Ocean microbes are key players in biogeochemical cycles that govern global ecosystem services [1, 2]. The pelagic zone, the largest habitat on Earth, is pervasively populated by picoplankton— organisms ranging from 0.2 to 3 *µ*m in diameter, including Bacteria, Archaea, and some viruses— that form a significant portion of the marine biosphere [3, 4]. These organisms are crucial for primary productivity, carbon and nitrogen fixation, and nutrient cycling, laying the foundation for the ocean’s trophic networks and supporting life across marine ecosystems. It is well-established that marine plankton communities are intrinsically linked to environmental conditions, but our knowledge of how the distribution of picoplankton communities relates to long-term climatic conditions is still poorly characterised. Although such communities are affected by both diel and seasonal variations in environmental conditions – mainly temperature and sunlight availability, especially at higher latitudes [5] –, genomic data has confirmed the existence basin-scale patterns of distribution in picoplankton communities that are stable year-round, superseding localised, seasonal, and short-term effects [6–11]. These biogeographical patterns coincide with oceanographic conditions such as the physicochemical composition of water masses and areas under the influence of major ocean currents, and vary depending on the plankton size fraction [7]. There are a few different models that propose a geographical partitioning of the oceans based on such abiotic properties, most notably the Longhurst provinces, initially proposed in 1995 and updated in 2013 [12, 13]. Bringing biology into such models, through the means of integration of molecular and climatic data, can offer crucial knowledge about the ecological role of picoplankton and its interaction with environmental factors [14, 15]. Establishing an evidence-based, interpretable, and reproducible model for the distribution of picoplankton communities is therefore essential for incorporating microbial observations into Earth System Models, and for clarifying the role of ocean microbes in the context of global change. Recent studies have identified biogeographical patterns in planktonic communities and used those to understand how projected future climate conditions may restructure the biogeography of planktonic organisms, predicting shifts in the distribution of zooplankton, phototrophs, diazotrophs, and subsequent effects in biogeochemical cycles [14, 15]. Such efforts, however, have been restricted to samples from one or two individual sampling consortia. There is therefore an urgent need for unified, integrated analyses in this field, to experiment with diverse approaches and methodologies, and to identify common trends. Here, we propose a global model of surface picoplankton biogeography, employing omics data integration methods to characterise it from multiple angles, and utilising 1454 samples from different sampling consortia. We present an in-depth description of the environmental, taxonomical, and functional composition of each biogeographical province. By utilising data integration approaches, our work proposes a new framework to obtain a deeper mechanistic understanding of plankton biogeography.

## Methods

An overview of our methodological procedure is shown on Supplementary Figure 1. Publicly available metagenomes (File S1 – Sample metadata, [16]) were mapped to a reference database of prokaryotic genomes to generate taxonomic and functional profiles (File S2 – Genome metadata, [16]). Pairwise metagenomic distance was measured between samples, and the resulting matrix was used to identify ten ecological groups using hierarchical clustering. Province area predictions were calculated by training a random forest classifier model with environmental data as predictors, and ecological group labels as the response variable. Multivariate analysis was used to identify environmental, taxonomic and functional features characterising each province.

### Metagenomic data collection

We compiled previously published datasets of ocean metagenomes that comprise data from TARA Oceans [17], GEOTRACES [18], Bio-GO-Ship and other smaller consortia [19, 20]. From those records, we removed the following types of samples: sediment (non-water), low oxygen, biofilm, samples from size fractions that were not prokaryote-enriched, and samples collected at a depth below 200m. We further removed 75 samples due to proximity with the coast or being contained within estuary or inland water bodies with no associated marine environmental data. This resulted in 2132 metagenomes across 1246 sampling stations (see File S1 – Sample metadata [16]). Following an iterative clustering procedure, we further removed samples below 25m depth (see Supplementary Material – Iterative Clustering Procedure), resulting in 1454 samples from 1122 sampling stations. These were collected between 77°S and 83°N of latitude and spanning 43 Longhurst provinces [12, 13]. Metagenomes were retrieved by their SRA accession numbers using the FetchNGS pipeline [21]. Multiple sequencing runs belonging to the same biological sample were pooled together for all downstream analyses.

### Environmental data collection

Environmental data was obtained from the Bio-ORACLE v3 server [22]. We downloaded global data layers for ten variables: ocean temperature, dissolved molecular oxygen, nitrate, phosphate, chlorophyll, salinity, silicate, pH, dissolved iron, and sea ice cover. We selected layers associated with surface waters and present time conditions and assigned environmental parameters to each sampling station by using the mean value of each parameter in that given geographical coordinate.

### Generation of reference database

To generate a standardised reference database for metagenomic profiling, we used Struo2 [23] to process a total of 15,551 genomes from the Unified Genome Catalogue of Marine Prokaryotes (UGCMP, File S2 – Genome metadata) [19], which uses standardised GTDB taxonomy [24]. Such genomes comprise a mixture of high-quality metagenome-assembled genomes (MAGs), and isolates deposited in reference databases (MarRef/MarDB and Genbank) The resulting reference database comprised 33,888,038 sequences, that were functionally annotated with EggNOG 5.0 database and the EggNOG-mapper tool [25, 26], resulting in 5,781 annotations at the gene level (KEGG Orthology identifiers) and 160 annotations at the KEGG pathway level.

### Coverage profiling

To perform functional and taxonomic profiling of metagenomes, we used the files downloaded with FetchNGS and ran automated quality control and trimming with Fastp [27]. We then used a custom pipeline to map the reads, sort and index alignments, and calculate coverage of coding sequences [28–33]. We used the trimmed mean values of coverage to generate abundance tables, which were then normalised by Bayesian multiplicative replacement of zeros and by the center-log ratio (CLR) transform [34, 35], following best practices for dealing with gene abundance data [36, 37].

### Clustering procedure

We defined provinces based on pairwise metagenomic distances between samples, calculated using Sourmash v4.8.2 [38]. Following the rationale from Richter et al [7], we chose the metagenomic distance metric that is reference-free metric and can therefore be applied to distinct types of datasets, eg. different size fractions of plankton. This distance was highly correlated to both taxonomic and functional structure (Supplementary Figure 2), confirming its usefulness as a proxy to community composition. We used the “average”, or UPGMA, linkage method, as recommended for hierarchical clustering in the context of biogeographical surveys [7], combined with an iterative clustering procedure (detailed in Supplementary Material – Iterative clustering procedure, Supplementary Figure 3).

### Variable selection

A goal of our analysis was to identify the most relevant taxonomical, functional, and environmental attributes that characterise each province. We analysed the abundance tables at the genus and genome level (for taxonomy) and at the KEGG KO and KEGG Pathway level (for function), and the environmental parameters from Bio-ORACLE. We then performed feature selection with Sparse Partial Least Squares Discriminant Analysis (sPLS-DA [35, 39]) that is well-suited for high-dimensional data. In each sPLS-DA model, we specified a given dataset as predictor variables and the province labels as the outcome variable. We also integrated different types of measurements from the same set of samples with the block sPLS-DA method that generalises sPLS-DA for more than one data set [40]. We focused on the selection of taxonomical and functional features but included all environmental features in the block sPLS-DA model.

### Province area projections

To predict province areas across oceans and project them across the global coordinate grid, we trained a random forest (RF) classifier model using the environmental parameters of each sampling station as the predictor variables and the province labels as the outcome variable. RF has been shown to be highly performant for this type of task, outperforming other approaches such as logistic regression classifier, gradient boosting machine, and generalized additive models [7, 14]. After performing model validation (see “Model validation”), we trained a RF classifier using the full dataset of 1122 sampling stations and fitted it with the environmental parameters of the full global coordinate grid of 16 million points. Hyperparameters for RF classifier are shown on Supplementary Figure 4A. To represent areas of transition between provinces, we decreased the opacity of data points where the confidence (ie. the fraction of decision trees in the forest voting for that province) in the predicted label was less than 0.75.

### Model validation

To test the validity and performance of the RF classifier, prior to training on the full dataset, we performed repeated stratified k-fold cross-validation. We averaged the results across all repeated folds to build a confusion matrix (Supplementary Figure 4A); we also constructed Receiver Operating Characteristic (ROC) curves from One-versus-Rest classifiers to assess how each individual province could be distinguished (Supplementary Figure 4B). Training and test samples for constructing ROC curves including 50% of the total dataset, stratified on the outcome variable (province). Moreover, to obtain the importance of predictors for each individual class, we calculated the Pearson correlation between the different environmental predictors and calculated each feature relative importance (Gini metric) across the validation folds. This was done both for the general RF model used for province area projection, and for the individual One-versus-Rest classifiers between all province classes (Supplementary Figure 4B and C, respectively). The Gini metric is calculated based on how much a feature decreases the Gini impurity across all trees in the forest, or how often a randomly chosen element from the dataset would be incorrectly classified if the samples were randomly labelled.

## Results

### A global biogeography of surface picoplankton communities

We propose a global model of picoplankton biogeography consisting of ten biogeographical provinces, divided into three categories: polar (two provinces), tropical (three provinces) and temperate (four provinces), and the Baltic Sea as an outlier (Figure 1A). We found a robust link between metagenomic composition and environmental factors, which strengthens the argument for a global, basin-scale partitioning of the oceans into biogeographical provinces which supersede local and seasonal variations. The robustness of this relationship between the molecular and climate data is evidenced by the performance of the machine learning classifier used to predict province areas (Figure 1B) (mean accuracy 0.94 ± 0.02, Supplementary Figure 5). Given that province labels were defined on metagenomic data alone, and the classifier used environmental data as predictors, this demonstrates how the community status, as represented by metagenomic composition, can be inferred from the long-term environmental makeup. The model showed a clear partitioning of the oceans into latitudinal bands; the tropical provinces tropical low nutrient and high nutrient (respectively TRLO and TRHI) and subtropical gyres (TGYR) were concentrated around the Equator and the tropics (with TGYR expanding up to 35.88°N, Table 1). TRLO was the largest province, covering nearly 30% of the surface of the global oceans. TGYR can be interpreted as a transition zone between the TRLO province and the colder temperate Mediterranean-like (MTEM) and subtropical/temperate transition (STEM) provinces. The regions where TGYR is located is associated with subtropical oceanic gyres, especially the South Atlantic and North Atlantic Gyres. The TRHI province was restricted to the eastern portion of the tropical Pacific Ocean, except for small areas near the mouth of the Amazon River and off the western coast of Africa. It is distinguished by a higher nutrient content, especially phosphate and nitrate, and lower pH, in comparison to the TRLO province (Figure 2). This province can be associated with the upwelling influence of the Humboldt Current, which feeds nutrient-rich waters into the South Equatorial Current. The temperate category included four provinces. The STEM province establishes a transition between the tropical and temperate zones, and its location the North Atlantic is influenced by the Gulf Stream Current. It also appeared as a latitudinal band in the southern hemisphere, dividing the subtropical TGYR province and the oceanic temperate (OTEM) province. The STEM province is a sister group to the MTEM, or temperate Mediterranean-like province. This province showed a diverse geographical distribution, by occupying the majority of the Mediterranean Sea, but also regions of the South Atlantic and South Indian Oceans (in the Agulhas Current region), parts of the North Atlantic Gyre, a small region of the Pacific off the eastern coast of Japan, and the southern Australian coastline. Interestingly, the samples from the Mediterranean Sea formed a single clade in the dendrogram of the subtree that contains this province, suggesting increased connectivity and a degree of geographical isolation for samples in this region (File S3 – Complete dendrograms [16] and Supplementary Figure 6). The OTEM province was largely comprised by a latitudinal band above the Southern Ocean, with its limit marked around the 60°S of latitude. It also appeared in the North Pacific and North Atlantic Ocean, as the province marking the transition between the warmer, more tropical TRLO/TGYR zones and the colder waters of the Boreal/Arctic province. The fourth province in the temperate category was the coastal temperate (CTEM), characterised by its proximity to coastlines, especially along the coasts of North and South America, Northern Europe, and the Yellow Sea. It marked the transition between the Baltic Sea province (BALT), and the adjacent OTEM; the Black Sea was also attributed to this province, although there are no sampling stations for this region in our model. It can be broadly described as a temperate coastal province that is correlated with areas under the influence of upwelling. The polar category consisted of the Antarctic province (APLR) in the Southern Ocean around Antarctica, and the Arctic/boreal province (BPLR) in the Arctic Ocean. Although the environmental makeup of both provinces was similar, the former displays slightly lower mean temperature, decreased sea ice cover and dissolved molecular oxygen, and a higher concentration of phosphate, nitrate, and silicate (Figure 2). The BPLR province showed an increased latitudinal range when compared to APLR, approaching 40°N latitudes in the eastern coast of North America and northern Japan. Although we described this province as “Arctic” due to all of the sampling stations classified in that province being contained in that region, the predicted area for this region also comprises a thin band between the APLR province and OTEM, in the Southern Ocean around the high-fifties degrees of latitude; this suggests that the environmental composition of this region is more similar to the Arctic region than to the neighbouring APLR province. However, it may be an artifact of the prediction model as there are no biological samples from that area. Lastly, the Baltic Sea province, which was the most divergent from other provinces (Figure 1A), displays far lower salinity (likely due to the large freshwater input in this region), and high mean concentration of dissolved iron and molecular oxygen. Identifying this province as an outlier is an important step to establishing a global model of picoplankton biogeography, as it suggests that other enclosed or semi-enclosed seas may display similar patterns, although a lack of representative sampling currently prevents this from being investigated. We have also observed a pattern of connectivity consistent with the latitudinal partitioning; the Arctic/boreal province was the most highly connected, that is, it showed the highest degrees of similarities between samples, followed by the Antarctic province (Supplementary Figure 6). While the former displayed some connectivity with neighbouring provinces, the latter was mostly disconnected. We also observed sampling stations that served as “hubs”, connecting geographically distant locations, such as the MTEM samples in the south-eastern and northern parts of the Atlantic Ocean.

**Figure 1.**
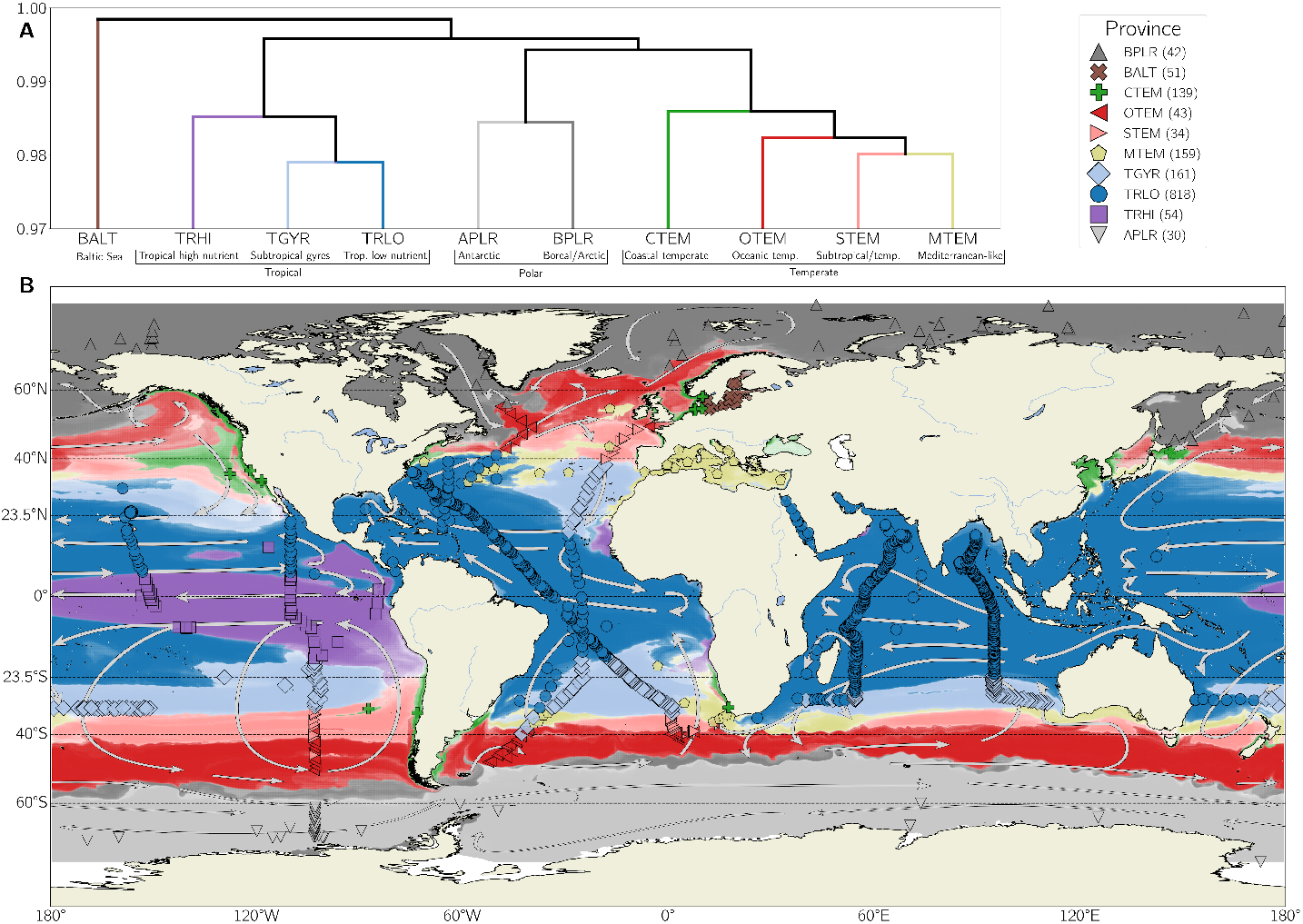
A global model of picoplankton biogeography. A) Collapsed dendrogram shows ten distinct provinces, divided into three categories (tropical, polar, temperate) and an outgroup. World atlas (B) shows the locations of sampling stations as coloured markers, with each colour and shape combination corresponding to a province (group and number of samples per group indicated in the top right legend). Shaded areas on the map show province area projections, based on a random forest classifier model trained by using the environmental parameters at each sampling station as the predictor variables, and the province label as the outcome variable. Arrows denote major ocean currents.

**Figure 2.**
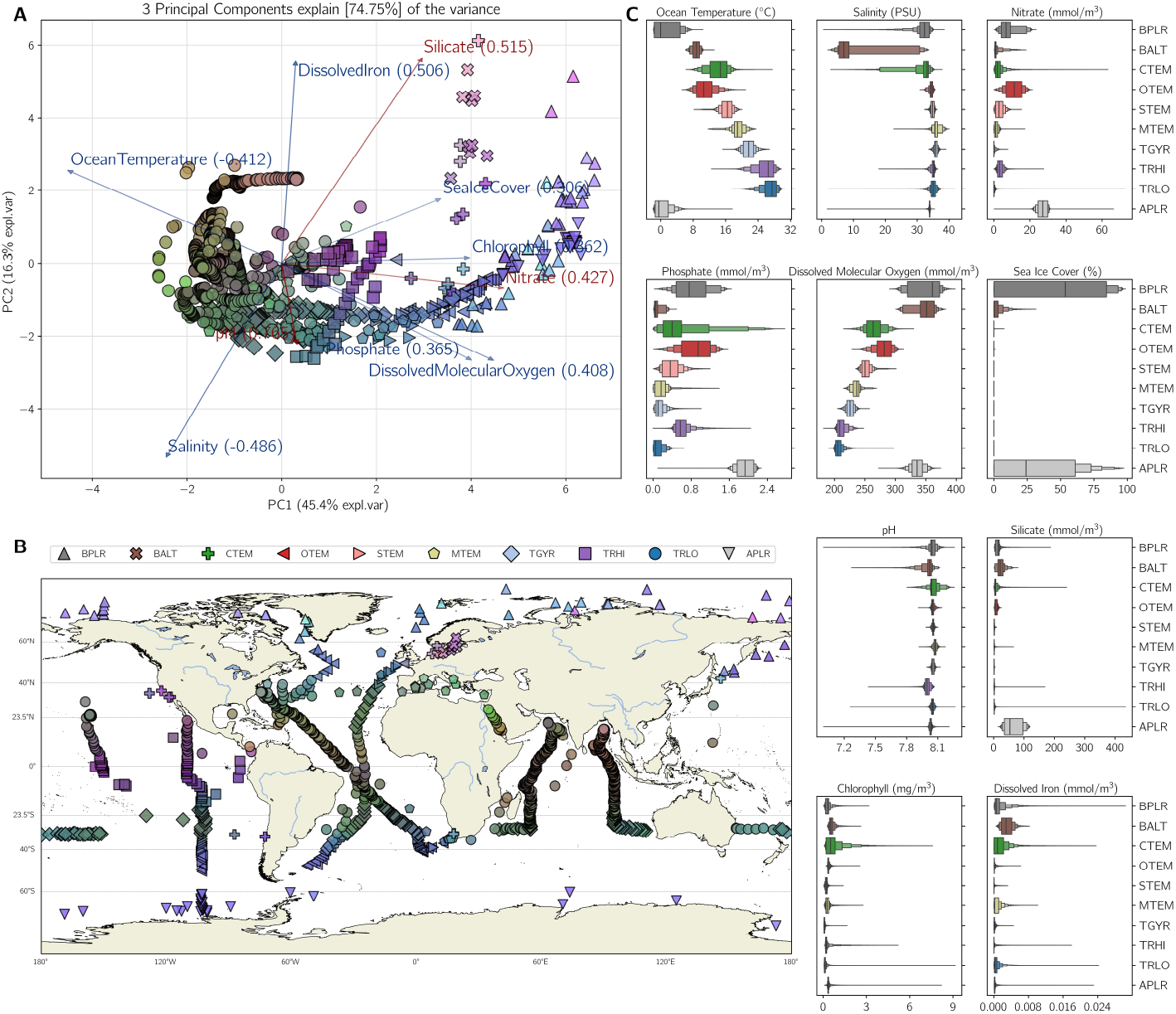
Environmental profiling shows a gradient consistent with community structure. A) Biplot of Principal Component Analysis (PCA) of ten environmental variables projected into three principal components (PC), including variable loading vectors displayed as arrows. Markers are coloured according to their position in the PCA, with each PC being mapped to a colour channel. B) world map of sampling stations. world map. C) letter-value plots of environmental variables across the different provinces, using environmental parameters for all coordinate locations classified as a given province (depicted in Figure 1B).

**Table 1.** Description of provinces. Provinces are labelled with a four-letter code to facilitate visualization, and are divided into four categories: polar, temperate, tropical, and Baltic Sea. Projected latitude and area correspond, respectively, to the range of latitude, and relative proportion of area, calculated in province area projections depicted in Figure 1. ‘No. S’ denotes number of samples in each province, and ‘Prev.’ denotes closest corresponding province/ecological status in previous classifications (province starting with “B” from [7] and with “ES” from [15]).

| Province | Description | Category | Projected Latitude* | Proj. Area | No. S. | Prev. |
| --- | --- | --- | --- | --- | --- | --- |
| **BPLR** | Arctic/boreal polar | Polar | 53.99°N – 85.00°N | 15.66% | 42 | ES2, ES5 |
| **BALT** | Baltic Sea | – | 44.89°N – 65.84°N | 0.18% | 51 | ES2 |
| **CTEM** | Coastal temperate/upwelling | Temperate | 19.98°N – 53.49°N<br>12.43°S – 49.34°S | 2.14% | 139 | ES5 |
| **OTEM** | Oceanic temperate | Temperate | 29.33°N – 63.54°N<br>21.38°S – 52.14°S | 12.73% | 43 | B6, ES6, ES3 |
| **STEM** | Subtropical-temperate transition | Temperate | 24.23°N – 47.79°N<br>21.13°S – 40.89°S | 5.15% | 34 | B7, ES6 |
| **MTEM** | Mediterranean-like | Temperate | 19.28°N – 42.18°N<br>11.79°S – 38.54°S | 2.99% | 159 | ES1 |
| **TGYR** | Subtropical oceanic gyres | Tropical | 14.98°N – 35.88°N<br>10.28°S – 32.88°S | 8.51% | 161 | B2, ES1 |
| **TRHI** | Tropical high nutrient | Tropical | 16.98°S – 12.53°N | 5.60% | 54 | B3, ES3 |
| **TRLO** | Tropical low nutrient | Tropical | 25.08°S – 29.18°N | 29.59% | 818 | B5, ES4 |
| **APLR** | Antarctic polar | Polar | 52.34°S – 76.95°S | 16.43% | 30 | B8, ES7 |

\*Projected latitude shows the values between the 0.1 and 0.9 percentiles of the absolute latitude range of the province, except for the BPLR, BALT, and APLR provinces, which show the values between the 0.1 and 1 values.

### Environmental profiling shows a gradient consistent with province area transitions

Although provinces were classified as discrete groups, environmental makeup at each sampling station showed a gradient that illustrates the rate of change in environmental parameters across geographical distance, and how each province transitions to another (Figure 2A and B). The environmental parameters that explain most of the variance between provinces are ocean temperature, nitrate, phosphate, dissolved molecular oxygen, and chlorophyll, the first of which is inversely correlated with ordination along Principal Component 1 in Figure 2A. Ordination along Principal Component 2 shows correlation with silicate, dissolved iron and, to an extent, ocean temperature, and inverse correlation with salinity, as shown by the Baltic Sea and Arctic Ocean samples in the top right corner of Figure 2A. Samples in the tropical Atlantic and in the Indian Ocean had fairly similar environmental makeup to the rest of the TRLO province, although transition to TGYR happened at a lower latitude in the southern Atlantic, and samples in the Bay of Bengal showed a distinctive profile with higher dissolved iron. The Red Sea, also classified as TRLO, also displayed a distinct environmental profile, with higher temperature and salinity when compared to other TRLO stations. We observed a sharp increase in nitrate and chlorophyll content in the equatorial portion of the eastern Pacific, marking the transition from TRLO to TRHI (Figure 2B). The temperate provinces MTEM, STEM and OTEM followed a gradient of increasing levels of phosphate, nitrate, dissolved molecular oxygen, and decreasing temperature, while CTEM stood out from its temperate counterparts due to increased chlorophyll content (Figure 2C). Although mean salinity was mostly uniform across provinces (except for the Baltic Sea), some provinces showed regions with decreased salinity (Figure 2C). Temperature was the most significant predictor to identify community composition, followed by dissolved molecular oxygen (inversely correlated with temperature, Supplementary Figure 5A), and then nitrate and phosphate (Supplementary Figure 5B). The latter two indicated differences between closely related provinces, and were increased in the tropical high nutrient, oceanic temperate and Antarctic provinces, when compared to other provinces in the same category. When classifying individual provinces versus the rest, the most important predictors for the Antarctic province were nitrate, phosphate and silicate, presumably what differs it from the Arctic/boreal province. For the tropical high nutrient, the most important predictor was pH, followed by temperature and phosphate. For subtropical gyres, it was chlorophyll, followed by temperature and dissolved molecular oxygen. For the Mediterranean-like, the key predictor was salinity, also followed by temperature and oxygen For the Baltic Sea and coastal temperate, the most important predictors were, by far, salinity and chlorophyll respectively, indicating that these parameters differentiate these provinces from their surrounding areas.

### Provinces display distinct community structure and dominant taxa

At the phylum level (Figure 3A), the tropical provinces showed the well-described pattern of high abundance of picocyanobacteria and small-celled heterotrophs, e.g. *Prochlorococcus* and *Pelagibacter*. Although both genera are ubiquitous throughout the oceans, they dominate surface communities in tropical waters [3]. Subtropical gyres (TGYR) showed an increased abundance of SAR86 and *Roseibacillus*, as well as the undescribed Gammaproteobacteria bacterium GCA-002705445 when compared to the tropical high nutrient (TRHI) and low nutrient (TRLO) (Figure 3C). TRHI showed an increased abundance of *Oceanicoccus*, the Dehalococcoidia bacterium UBA1127 and the Pseudomonadales bacterium HTCC2089. TRHI also showed a higher median relative abundance of Archaea when compared to TGYR and TRLO. The most abundant archaeal genomes in the TRHI province were MG-IIa (Marine Group IIa) and *Poseiidonia* genera [41]. TGYR and TRLO also showed high prevalence of the archaeal group MG-IIa, but the other most abundant genome belongs in the order MG-III for the former and of the family Thalassarchaeaceae (MG-IIb) for the latter.

**Figure 3.**
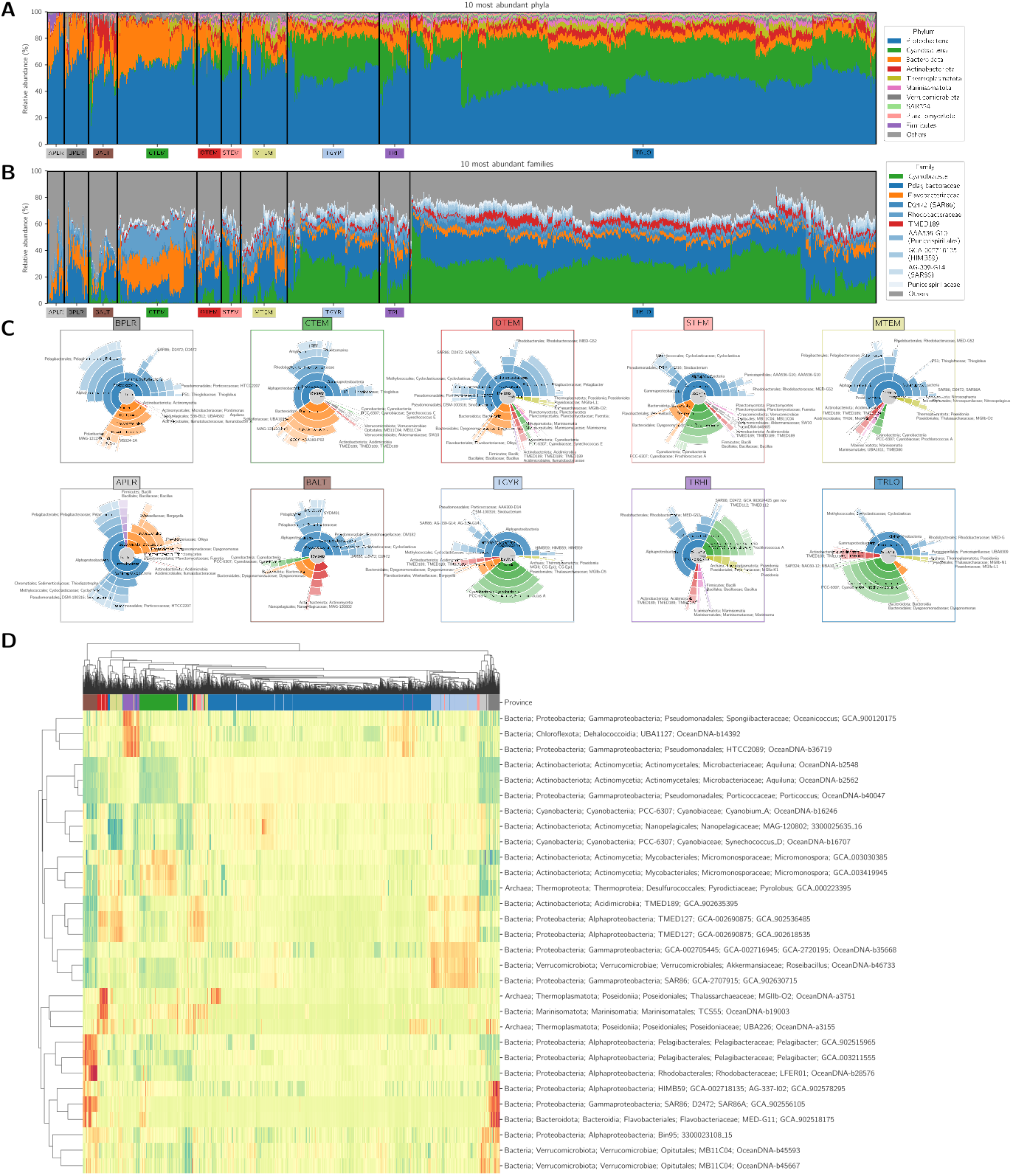
Taxonomic profiling shows distinct dominant taxa across provinces. Top: relative abundance of the ten most abundant phyla (A) and families (B) across all samples, as per highest median value. C) representative communities of provinces (see Supplementary Material – Representative communities of provinces). Sunburst chart shows the relative abundance of taxonomical ranks that each genome belongs to, from domain (innermost circle) through phylum, class, order, family, and genus. Bottom: Cluster image map of the features selected by sPLS-DA, showing correlation between genomes (rows) and samples (columns). Rows and columns in the heatmap are ordered according to UPGMA clustering, with corresponding dendrograms.

The transition from tropical to temperate provinces is accompanied by a decrease in the relative abundance of cyanobacteria. Temperate provinces also showed a remarkable increase in Bacteroidota when compared to tropical provinces. This is especially true for the CTEM province, where there was a high abundance of Flavobacteriaceae and Rhodobacteraceae (Figure 3B). At the phylum level, the polar provinces showed similar community structure to coastal temperate and oceanic temperate, although the Antarctic (APLR) also shows increased abundance of Firmicutes. This did not hold at the family level, where the most abundant families across all samples comprised a smaller fraction of the total relative abundance in the polar and Baltic provinces. Representative communities for each province (Figure 3C) described the difference in abundance between dominant genomes in each community. For APLR, although the median relative abundance of Archaea was only 0.4%, a genome of the methanogen *Methanococcus maripaludis* (Archaea, Methanobacteriota, accession GCA-002945325) was amongst the ten most abundant. APLR also showed remarkable diversity of families in the Gammaproteobacteria class; the genus Cycloclasticus, which degrades aromatic compounds, appeared with high relative abundance (median of 4.2%), as well as other metabolically versatile genus such as *Thiodiazotropha* (a symbiont with the capacity to fix nitrogen), *Sinobacterium* (moderately halophilic nitrate reducer), and HTCC2207/SAR92 (DMSP degrader) [42–45]. APLR also showed a high abundance of Bacillus subtilis (2.8%) and of genomes in different families in the order Bacteroidales, such as Flavobacteriaceae, Weeksellaceae, and Dysgonomonadaceae. At the phylum level, Boreal/Arctic (BPLR) showed similar structure to APLR, although without the high abundance of Firmicutes. This was also indicated by the lack of a genome in this phylum (such as *Bacillus subtilis*) in the representative community. BPLR displayed high abundance of common heterotrophs such as *Pelagibacter* (Alphaproteobacteria), SAR86, and HIMB59 (both Gammaproteobacteria) (Figure 3D). Samples from the Baltic province stood out at the phylum level for having a high relative abundance of Actinobacteriota and Planctomycetota. This province displayed a dominance of bacteria of the families Nanopelagicaceae, Microbacteriaceae (genus *Aquiluna*) and of the order Methylacidiphilales (Verrucomicrobiota) (Supplementary Figure 7). The representative community of the Baltic Sea also showed the presence of the genus *Cyanobium*_A (commonly misclassified as freshwater *Synechococcus*, [46]) (Supplementary Figure 8A). A few genera also presented genotype abundance patterns that reflected the biogeographic partitioning. For example, when looking at different genotypes of *Prochlorococcus*, we observed high abundance of *Prochlorococcus*_A (high-light group) in the tropical provinces, and *Prochlorococcus*_B (low-light group) in the temperate and polar provinces (Supplementary Figure 8B), consistent with previous observations of distributions of different *Prochlorococcus* genotypes [47]. We also found such patterns for individual *Prochlorococcus* genomes; for example, the tropical high nutrient province showed a high abundance of an individual genome of *Prochlorococcus* when contrasted with tropical low nutrient (Supplementary Figure 8D and F), and for heterotrophic bacteria in the families Pelagibacteraceae, SAR86, and HIMB59 (Supplementary Figure 8C, E, and G).

## Functional profiling reveals metabolic and ecological trends

The sPLS-DA analysis carried out to identify the most relevant features in the functional profile of each province showed different trends for each province category. For tropical provinces, there were increased concentrations of the photosynthesis pathway, as well as the biosynthesis of N-Glycan, carotenoids, terpenoids, and steroids, and the metabolism of porphyrin, lipoic acid, sugars (starch, sucrose, fructose, mannose), biotin, and thiamine. The Baltic province showed enriched pathways for biosynthesis of macrolides and different types of O-glycan. Features selected on the second component mostly characterised the APLR province, including the metabolism of retinol, xenobiotics, beta-Alanine, and amino acids (tyrosine, arginine and proline), degradation pathways for chloroalkanes and chloroalkenes, fatty acids, naphthalene, limonene, branched amino acids (valine, leucine, iso leucine), and styrene. Interestingly, a pathway maximised for that same province is that of insect hormone biosynthesis; although there are reports of the influence of the insect microbiome in hormone production [48] and of such biochemical compounds in aquatic habitats, its function in the planktonic ecosystem is largely unknown. Features selected on the third component also display interesting pathways which are maximised in the APLR, and partly in the Baltic, provinces: the degradation of toluene, fluorobenzoate, chlorocyclohexane and chlorobenzene, and biosynthesis of unsaturated fatty acids, arginine, and the biosynthesis of monobactam antimicrobials. Other antimicrobials, such as tetracycline, were maximised in the Mediterranean-like Temperate province. Features selected on the fourth component highlight antimicrobial pathways such as the biosynthesis of vancomycin and streptomycin (maximised in the tropical low nutrient province), resistance to vancomycin (maximised in the Antarctic province), and biosynthesis of acarbose and validamycin (maximised in the coastal temperate province). The Arctic/boreal, oceanic temperate and subtropical/temperate transition only present a single pathway that is maximised under our feature selection model, respectively the biosyn-thesis of siderophore group nonribosomal peptides, responsible for iron scavenging; the metabolism of alpha-Linolenic acid; and the biosynthesis of type I polyketide structures, a class of compounds that presents antibiotic properties [49].

### Metagenomic data integration untangles differences between closely related provinces

We next investigated the underlying differences between closely related provinces, ie. differences within the polar, temperate, and tropical province categories, by means of metagenomic and environmental data integration (Figure 5, Supplementary Figure 9, Supplementary Figure 10). To use the polar category as an example, the Antarctic province showed increased presence of the family Rhodobacteraceae (Alphaproteobacteria), in metabolism of drugs and xenobiotics, fatty acid degradation, and porphyrin metabolism, as well as increased phosphate and nitrogen content; markers for the Arctic/boreal samples were taxa the family Pseudohongiellaceae (Gammaproteobacteria), bisphenol degradation pathways, as well as increased pH, dissolved iron, and dissolved molecular oxygen. Some features could be attributed to variation within samples of the same province, such as vancomycin resistance, ubiquinone and zeatin biosynthesis, and taxa of the family Nitrincolaceae (Gammaproteobacteria), as well as environmental parameters of ocean temperature, salinity, chlorophyll, and sea ice cover (Figure 5A and B).

## Discussion

We present a model of picoplankton biogeography with unprecedented scale and resolution, utilising data from multiple sampling consortia and integrating different blocks of metagenomic and climatic data. Our results confirm and extend findings of previous studies that a large-scale partitioning of microbial communities in the oceans can be inferred from metagenomic data [7, 8, 14, 15, 50]. We have demonstrated how environmental data can be used to classify geographical areas into biogeographical provinces, and described the relationships between the taxonomic, functional and environmental features at each location, identifying features characterising each province.

### Community structure can be accurately predicted from environmental data

Our results demonstrate that picoplankton community structure can be accurately predicted by environmental parameters, showing the link between physical and chemical conditions on microbial biogeography (Supplementary Figure 4). We also showed that environmental data alone, when integrated with metagenomic information, can provide reliable insights into the distribution and diversity of microbial communities. This predictive capability underscores the importance of environmental drivers in structuring microbial biomes and offers a framework for forecasting shifts in microbial populations in response to climate change and anthropogenic disturbances.

### Undescribed taxa are major determinants of marine microbial biogeography

A notable aspect of our findings was that undescribed, uncultivated or generally poorly characterised taxa were major determinants of picoplankton biogeography. Recent advances in genome-resolved metagenomics and the generation of MAGs have implicated in hundreds of novel ocean microbial taxa being proposed through cultivation-free methods. An example of this is the gammaprotebacteria SAR86, first discovered over 30 years ago, but cultured only recently [51–53]. This taxon has displayed global biogeographical patterns correspondent to lineage-level genomic differences, ie. different lineages possess distinct gene clusters that affect its fitness in relation to environmental factors, and thus its large-scale distribution [54]. This is shown here in a community context (Figure 3C, Supplementary Figure 8). Other examples include the alphaprotebacteria HIMB59, also with few cultured representatives and disputed phylogenetic placement [55], the actinobacteria TMED189 (proposed order name Candidatus Actinomarinales [56]), and archaea of the Marine Group II (MGII, proposed order name Ca. Poseidoniales [41]), all of which are highly abundant in provinces of the tropical category. All of these share small genomes, suggesting metabolic streamlining, a common evolutionary strategy in oligotrophic environments, operates in the tropical surface oceans [57]. A proposed hypothesis to explain genome reduction on ocean microorganisms is that of adaptive gene loss in exchange for relying on co-occurrence or interactions with other organisms in order to obtain “public goods” (the Black Queen Hypothesis, [58]). Thus, it is relevant to examine the distribution of these taxa in both the community and data integration (ie. along with functional and environmental measurements) contexts. Several publications have described the ecological importance and ubiquity of these organisms, but they remain with few or zero cultured representatives. This work further confirms such importance, their abundance in representative communities of each biogeographical province (Figure 3), and their presence as discriminant features between provinces (Figure 5, Supplementary Figure 9, Supplementary Figure 10).

Carbon cycle and antimicrobial pathways are correlated with biogeographical patterns A remarkable functional aspect of the ocean microbiome is the diversity of carbon consumption and degradation pathways; the global oceans are one of the largest reservoirs of organic matter in the planet. As such, the metabolic mechanisms that microbes use to consume these compounds directly influences the microbial carbon pump and, by consequence, the global carbon cycle [59]. Our results showed how differences in functional profiles between provinces are largely shaped by distinct carbon degradation and consumption pathways. This is shown by the increased concentration of photosynthesis genes in the tropical provinces, which indirectly supports the widely accepted hypothesis that primary productivity in the tropical surface oceans is dominated by cyanobacteria, but is gradually taken over by eukaryotic microalgae as the distance from the Equator increases [60, 61]. The relationship between carbon cycle genes and biogeography is also demonstrated by pathways responsible for breaking down aromatic compounds such as naphthalene, styrene, toluene, fluorobenzoate, and chlorobenzene; such pathways particularly significant in higher-latitude areas. The concentration of such compounds can be largely attributed to atmospheric deposition, supporting the impact of atmospheric processes in microbial carbon cycling in the oceans [62]. Another aspect of the functional analysis is the prominence of genes for both antimicrobial resistance (AMR) and biosynthesis of antimicrobials being important features to discriminate between provinces. While the aromatic compound degradation pathways were mostly found in higher latitudes, AMR genes were widespread across provinces. Such genes have been found to be enriched across multiple habitats, not only marine [63]. The biosynthetic potential of the ocean microbiome is vast, and its biogeographical structuring is still poorly understood. Our results confirm previously reported trends [64], such as the tropical oceans being enriched in the synthesis of terpenoids (that have significant biological activity), and colder, nutrient-rich regions having a higher concentration of genes related to the synthesis of polyketide structures and of nonribosomal peptides (Figure 4).

**Figure 4.**
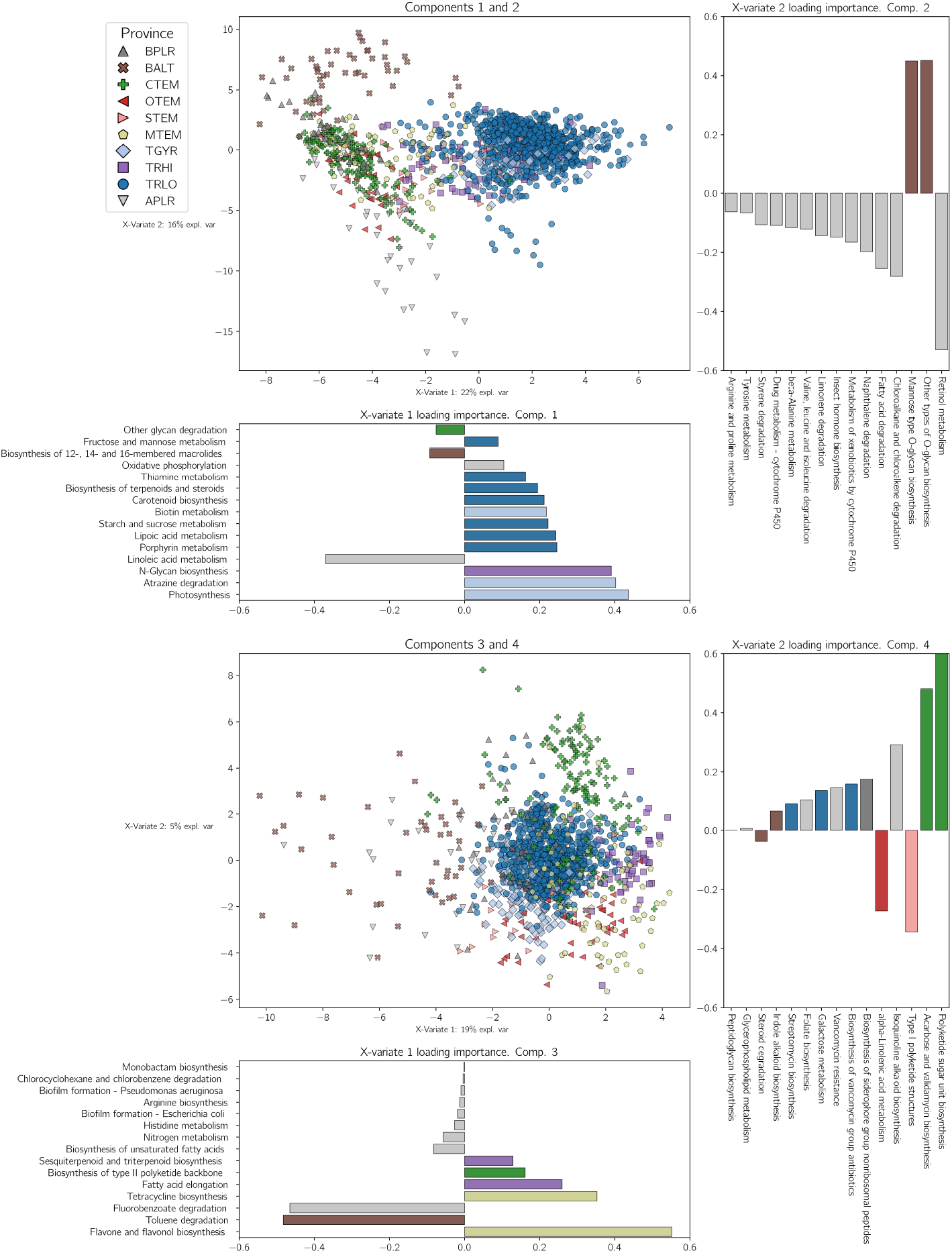
Picoplankton biogeographical provinces have distinct functional capacities. Ordination scatter plots of functional profiles based on KEGG Pathways, generated from sPLS-DA model with four components. The importance of features (loading coefficient) is represented on adjacent bar plots. Axes in the ordination plots show position of samples in the latent space, and degree of variability explained. Feature importance bar plots are coloured according to the class which has the maximised value for that given feature.

**Figure 5.**
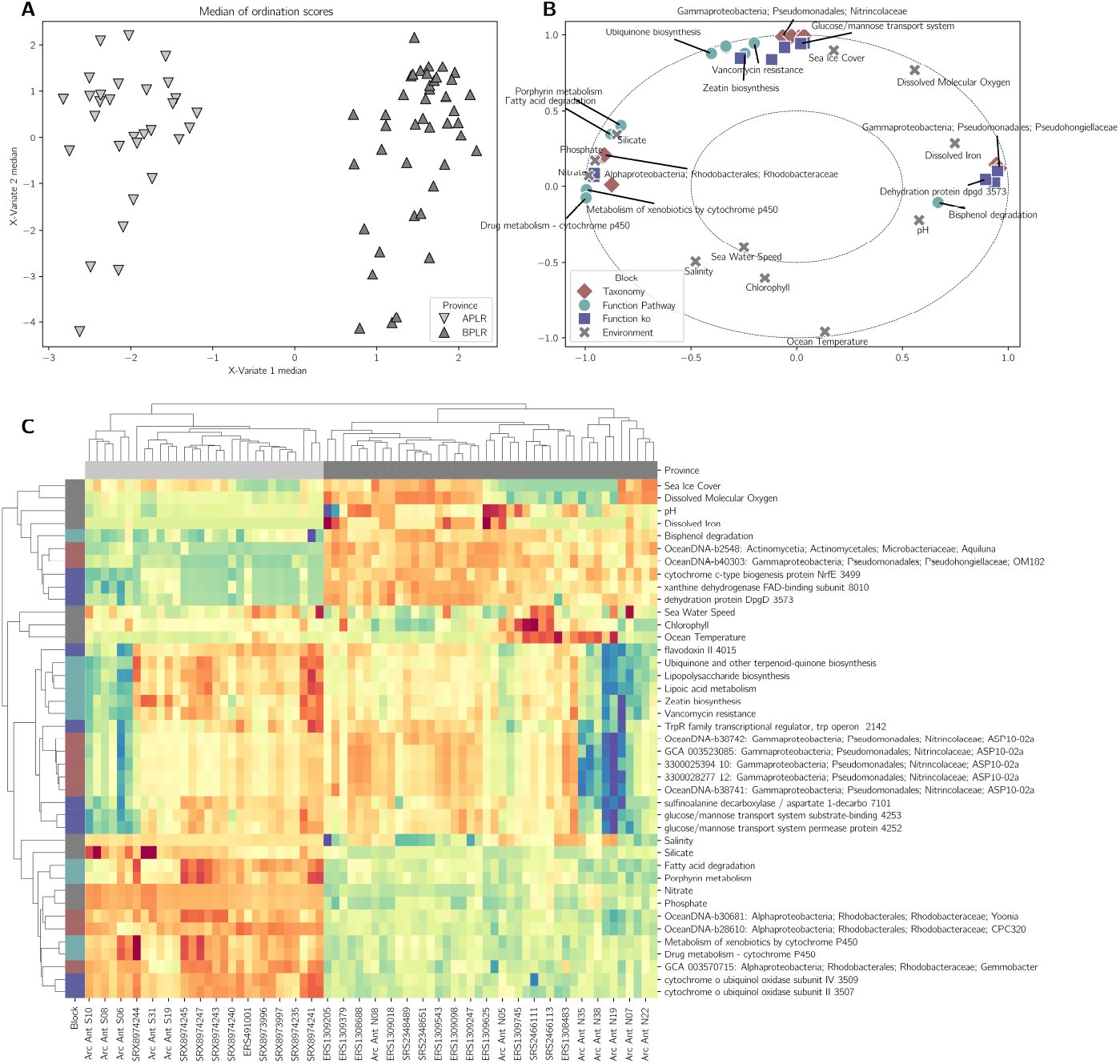
Metagenomic data integration highlights differences between polar provinces. A) median of ordination scores across the four datasets (coverage of genome, KEGG Pathway, and KEGG KO, and environmental parameters) from the multivariate integrative analysis with block sPLS-DA. B) Correlation circle plot displaying associations between features; position of features within the axes of this panel corresponds to their association with samples positioned along the axes in A. C) Clustered image map showing the association between samples and selected features.

### Microbial biogeography in the context of global change

Metagenomic data permits tapping into the genomic diversity of marine microbes, enabling insights into how it structures functional and taxonomic differences. Our study stresses the need to classify and describe biogeographical provinces as products between oceanographic features and genomic composition, a concept presented by Frémont et al [14] as climato-genomic provinces. Moreover, it is critical to adequately name such provinces to reflect their geographical location and/or their biotic and abiotic attributes, as done for the Longhurst provinces and proposed by Elizondo et al for phytoplankton groups [8]. A significant limitation of current models of marine microbial biogeography is the lack of data from multiple plankton size fractions from the same sampling locations; although biogeographical partitioning has been proposed for higher size fractions [7], uneven sampling creates a hurdle when integrating data from multiple fractions [65]. A promising approach is the usage of universal sampling protocols that target all three domains of life [66], and the standardisation of protocols for marine microbiome sampling across consortia. We argue that future research should move in the direction of conciliating different models of biogeography, consolidating common genomic and environmental drivers determining biogeographical patterns, and establishing consensus so that microbial and molecular observations can be effectively integrated into Earth System Models [67–69]. This will contribute to the robustness of both present-time and future climate scenarios and improve our understanding of the influence of ocean microbes on global ecology.

## Acknowledgements

VWS is funded by a Melbourne Research Scholarship from The University of Melbourne. HV is supported by a fellowship from the Fundação para a Ciência e a Tecnologia (CEECIND:2023.06155). VRM is supported by the Australian Research Council (DE220100965). KALC is supported by the by the National Health and Medical Research Council (NHMRC) Investigator Grant (GNT2025648). This research was supported by The University of Melbourne’s Research Computing Services and the Petascale Campus Initiative. We also thank the following individuals for insightful discussions: A/Prof Lucie Bittner, Marko Terzin, Dr. Saritha Kodikara, and members of the Lê Cao and Verbruggen labs.

## Data availability statement

The datasets analysed during the current study are available in the NCBI Sequence Read Archive, the Bio-ORACLE v3 server, and the OceanDNA Figshare record, https://www.ncbi.nlm.nih.gov/sra, https://erddap.bio-oracle.org/, https://doi.org/10.6084/m9.figshare.c.5564844.v1. Supplementary Material is available from UniMelb Figshare at https://doi.org/10.26188/27872490. Code used to process the data and generate figures is available from GitHub at https://github.com/vinisalazar/global_picoplankton_biogeography_2024/. Software used to process the data include libraries Cartopy, Dask, Geopandas, Matplotlib, Numpy, Pandas, Scikit-Learn, Scipy, Seaborn, Xarray, dplyr, mixOmics, stringr [33, 35, 70–77]. Software versions are available in the GitHub repository.

