## Supplementary Material for "Global picoplankton biogeography revealed by metagenomic and climatic data integration"

For the article "Global picoplankton biogeography revealed by metagenomic and climatic data integration", Salazar *et al.*, posted in November 2024.

---

#### Metagenomic distance

We calculated the metagenomic distance (or dissimilarity) between samples using Sourmash v4.8.2 [38]. We first calculated the k-mer signature of each metagenome individually post quality control, using the `sourmash signature` command, then generated a distance matrix using the `sourmash compare` command with the option `--scaled 1e6`. Similarly to the coverage data, we pooled SRA Runs with the same Sample accession number by using the 'sourmash signature merge' command.

#### Iterative clustering procedure

To define biologically and statistically cohesive provinces, we performed an iterative clustering procedure described as the following: i) generate of a linkage matrix from the distance matrix; ii) define clusters by cutting the linkage matrix into a predetermined k number of clusters; iii) remove of outliers, *ie.* clusters with less than 10 samples, but usually singletons; iv) remove outliers from linkage matrix; and repeat. At each round, if one cluster was much larger than the others, it would be analysed separately as a subtree of the linkage matrix, using the same procedure. We performed three rounds of clustering in total: in the first round, we used all 2132 samples, established $k = 8$ and removed three singletons, being left with five clusters. We then analysed the subtree of the largest province (containing 1482 samples). We set $k = 6$ for that subtree and removed three more singletons, dividing the subtree into three groups, totalling 7 clusters across all samples. After this initial round of clustering, we analysed sample collection depth between provinces and found that two provinces (2 and 21) stood out as having a far higher mean sample collection depth. This partitioning is illustrated in Supplementary Figure 3. After ascertaining this, we applied a more stringent depth cut-off and only used samples collected above 25 meters of depth, representing the very surface of the water column, so that depth would not be a confounding variable when defining clusters. In the second round of clustering, we used 1512 samples after filtering by depth. We set $k = 6$ and removed two more outliers, being left with 4 groups: Temperate, Polar, Tropical, and Baltic Sea provinces (Figure S6 – Collapsed dendrograms). We analysed the subtree of the Tropical province, the largest group with 1056 samples, setting $k = 14$, and removed 23 outliers, being left with 3 subgroups within that subtree. We performed a third round of clustering with the remaining 1487 samples, setting $k = 18$. This led to the removal of 33 additional outliers and resulted into a partitioning of 10 robust provinces across 1454 samples (Table 1).

#### Representative communities of provinces

By default, we selected the ten most abundant genomes by calculating the median relative abundance across samples in the province. For provinces that had at least 2% median relative abundance of Archaea, we also selected the two most abundant archaeal genomes. If the ten selected genomes belonged to three phyla or less, we selected an additional ten genomes from other phyla.

### Supplementary Figures

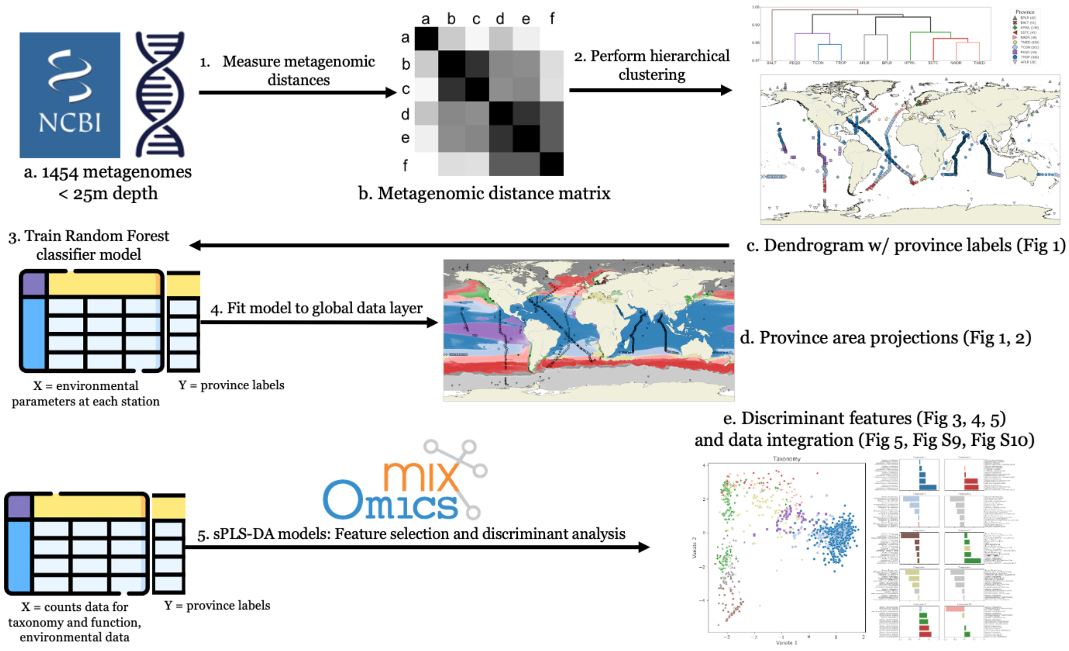


**Supplementary Figure 1. Methodology overview.** Diagram describing simplified methodology, with procedures (1 to 5) and results (a. to e.). Not shown are the coverage profiling procedure used to generate counts data for taxonomic and functional profiles, and random forest model validation.

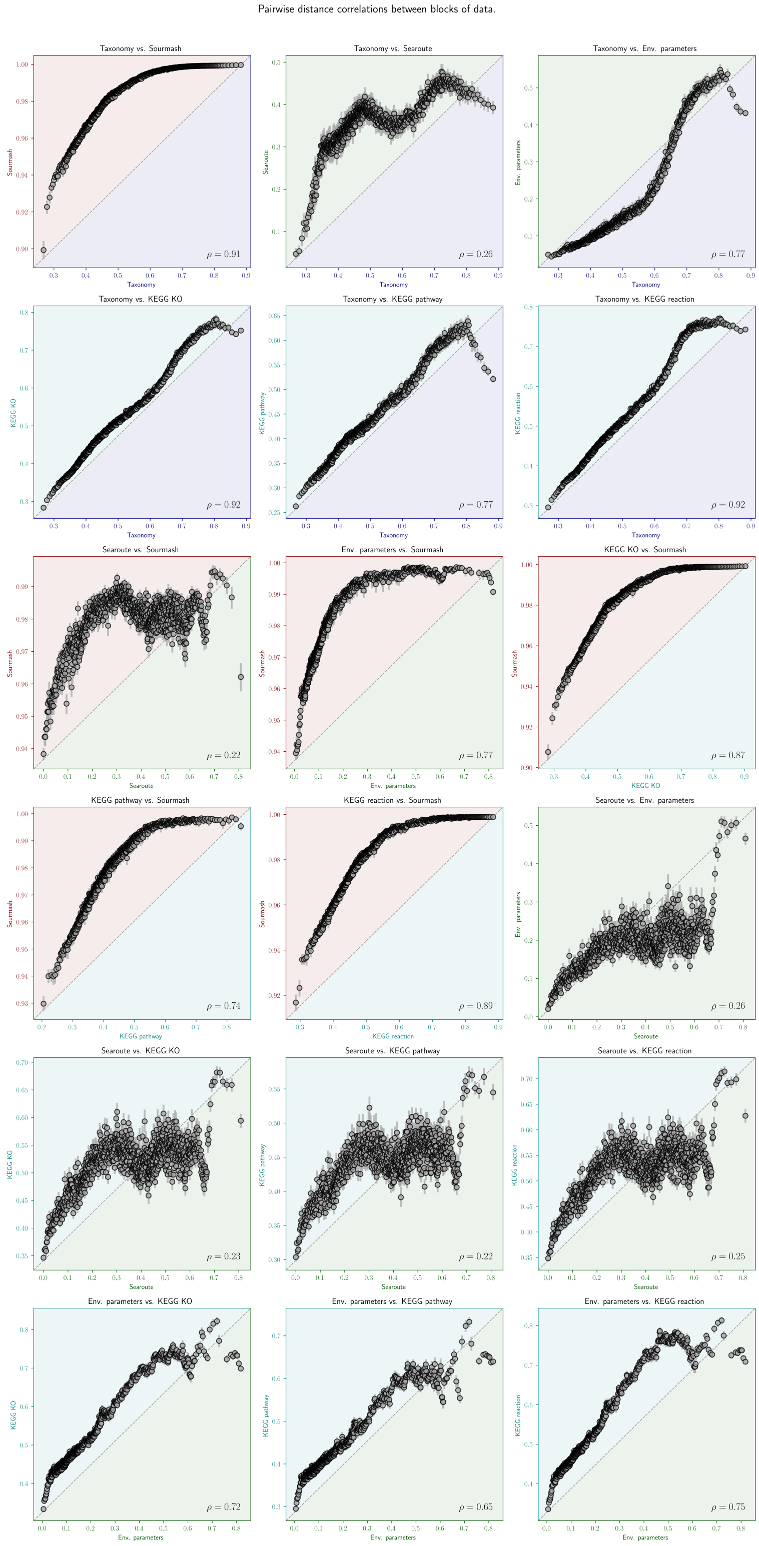


**Supplementary Figure 2. Distance correlations between blocks of data.** Different data blocks used for integrated analysis present varying degrees of correlation. Scatterplots show pairwise distance between samples across different data blocks (labelled on X and Y axes). Each point shows a bin of 1000 pairwise comparisons, with bars denoting the range of the bin. Correlation coefficient is shown on bottom right of each plot. Taxonomy (at the genome level) is shown in purple, k-mer level metagenomic distance in red, functional profiles in cyan, and environmental variables (including geographical distance) in green. We calculated the geographical distance between samples using the searoute v1.2.2 package [1].

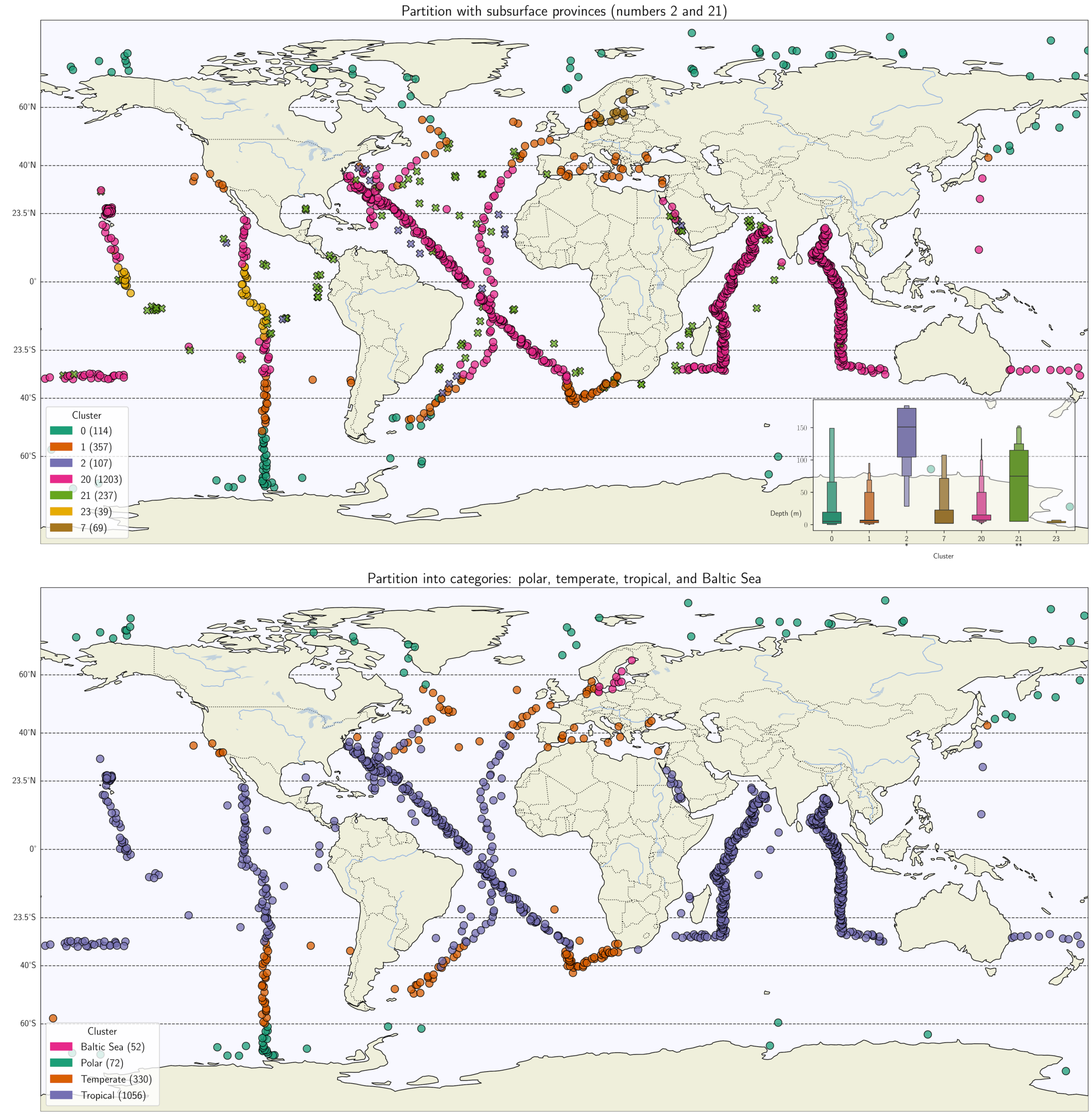


**Supplementary Figure 3. Subsurface and category-based biogeography models.** Before reaching our final model, we experimented with different interpretations of the hierarchical clustering dendrogram. Our first proposal (top) contained seven provinces, but we identified two of these provinces (2 and 21) as being at higher depths than the other five. Distribution of depth of sampling stations across clusters is shown on the inset. Asterisks under cluster names on the X axis indicate significant differences with other groups (Pairwise Tukey HSD test). This suggests the existence of a distinct subsurface biogeography (see [2] for discussion). After this, we removed all samples beneath the depth of 25 meters. This led to the second proposal (bottom), where the hierarchical clustering dendrogram is divided into four broad categories: polar, temperate, tropical and Baltic Sea. We did a further round of iterative clustering to remove singletons and reach the final model displayed in Fig 1 (see ).

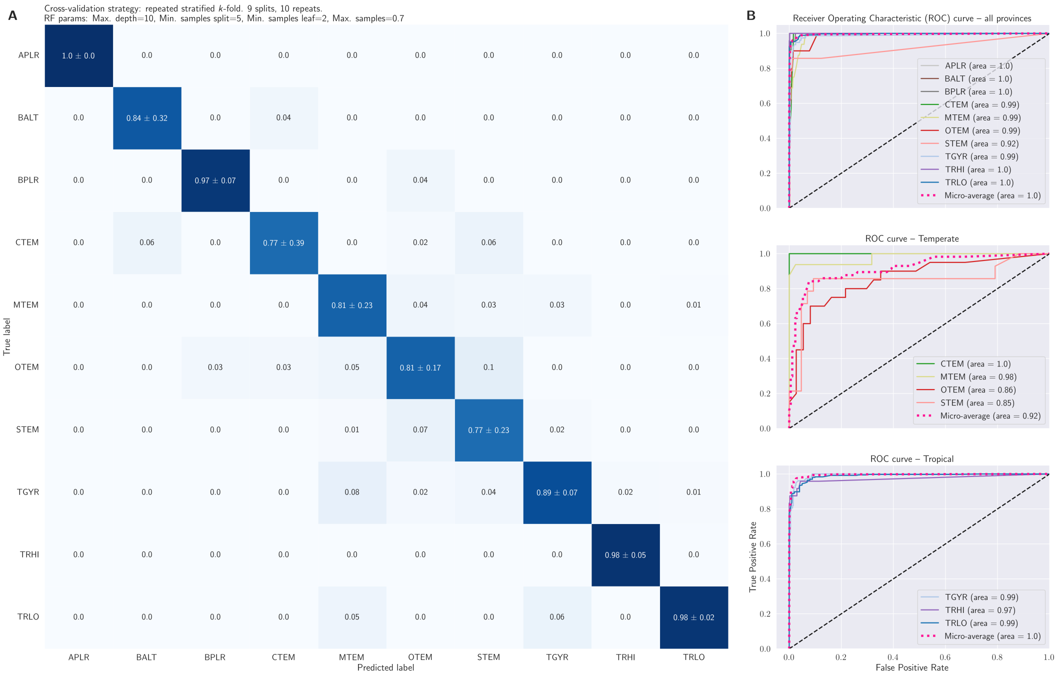


**Supplementary Figure 4. Confusion matrix and ROC curves of classifier models.** Panel A: Confusion matrix of repeated stratified k-fold cross-validation (CV) of random forest (RF) model. Model and CV parameters are shown on the top left. Numbers within tiles of confusion matrix indicate mean proportion of true and selected label across rounds of CV, plus and minus the standard deviation. Panel B: Receiver Operating Characteristic (ROC) curve for a one-vs-rest RF classifier between provinces. From top to bottom: between all provinces, between temperate provinces, and between tropical provinces. ROC curve for binary classifier between polar provinces was omitted as the model displayed perfect performance.

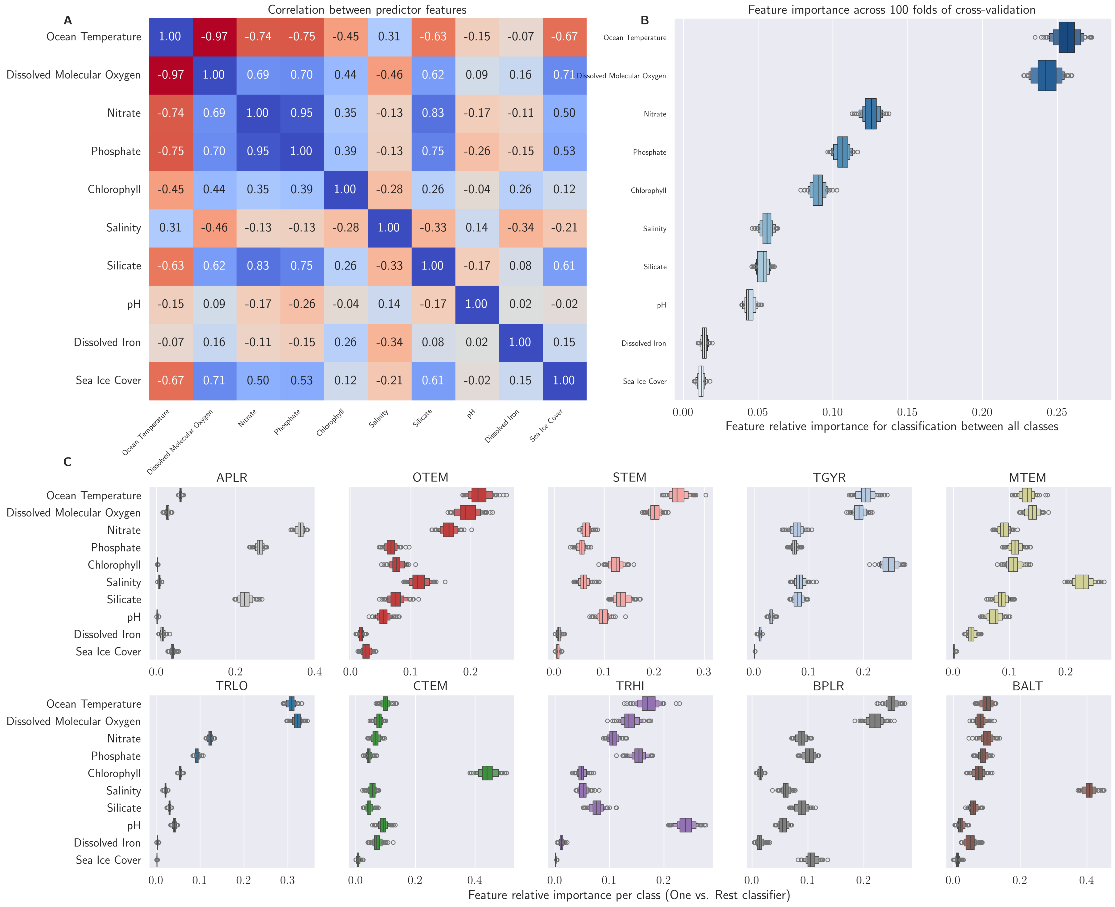


**Supplementary Figure 5. Feature importance and correlation.** Feature importance of environmental predictors for classification of province labels. A shows correlation between predictors in the training dataset. B shows mean relative feature importance for general random forest classifier across 100 folds of cross-validation. C also shows the mean relative feature importance for 100 folds of CV, but for individual estimators of each class in a One versus Rest classifier.

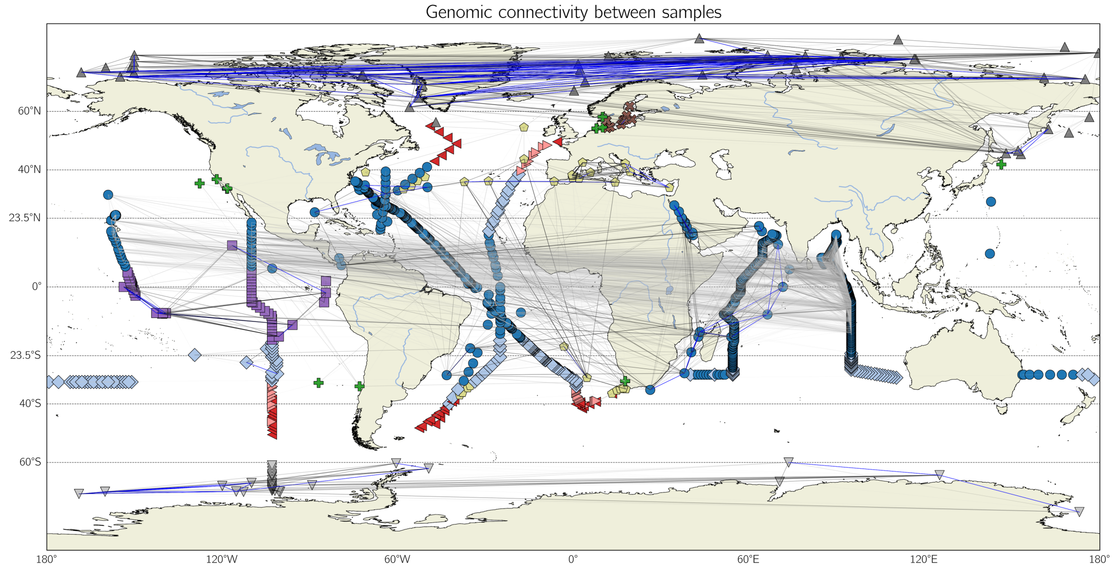


**Supplementary Figure 6. Map of global sample connectivity.** World map with sampling stations (coloured according to proposed province), with links denoting degree of metagenomic connectivity. Light gray links indicate metagenomic similarity values above the 0.5 quantile, dark gray, above the 0.75 quantile, and blue links, on the upper 0.9 quantile, considering all pairwise metagenomic distance comparisons between sampling stations. Sampling stations are coloured according to provinces, as in Figure 1.

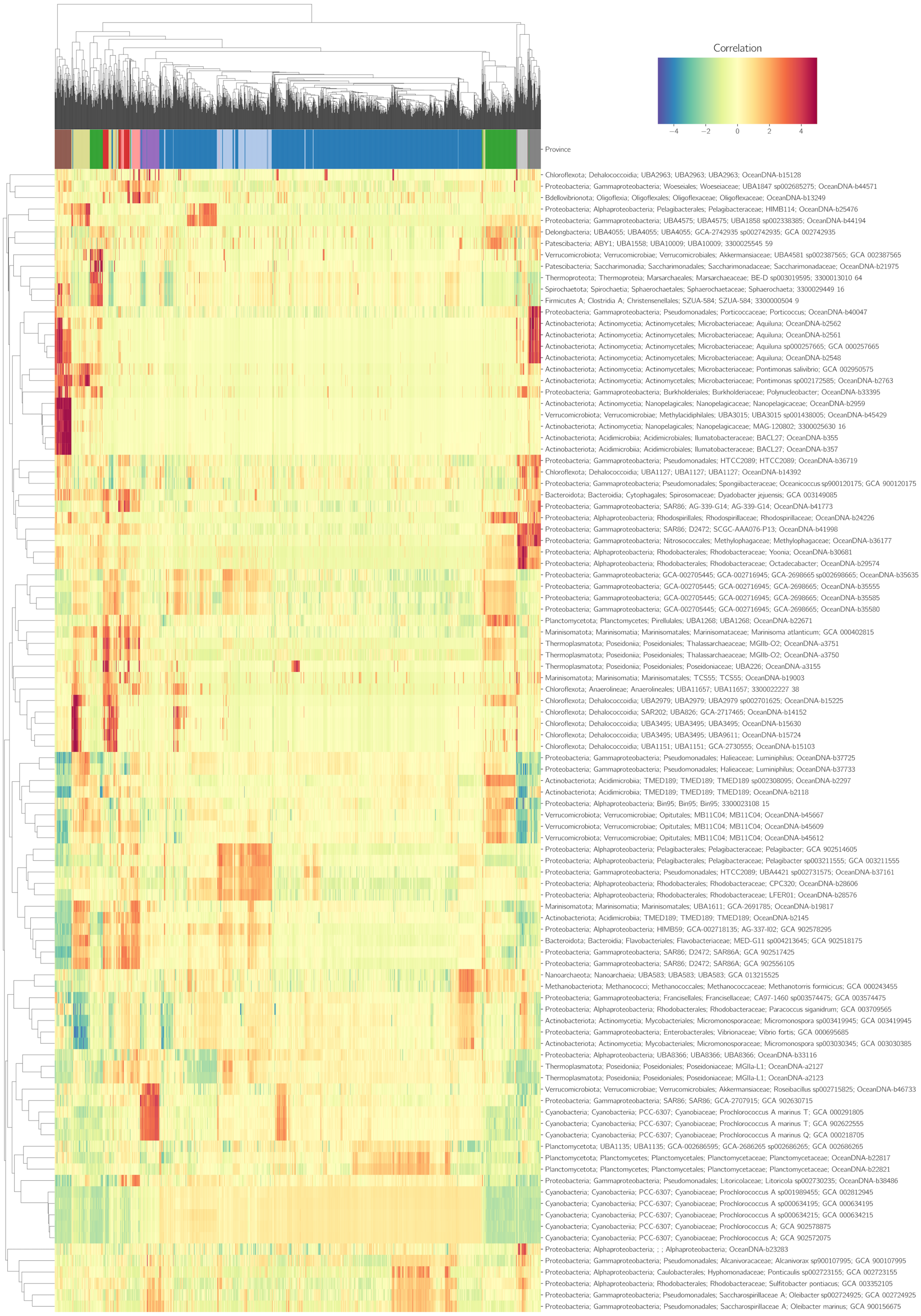


**Supplementary Figure 7. Expanded cluster image map of genome-based sPLS-DA model.** Expanded cluster image map for sPLS-DA model for feature selection and classification. Model was built with 5 components and 20 features per component. Rows indicate features, and columns indicate samples, with colour indicating degree of correlation between features and samples. Topmost row indicates province label of each sample. Dendrograms on X- and Y-axis show hierarchical clustering, average/UPGMA method.

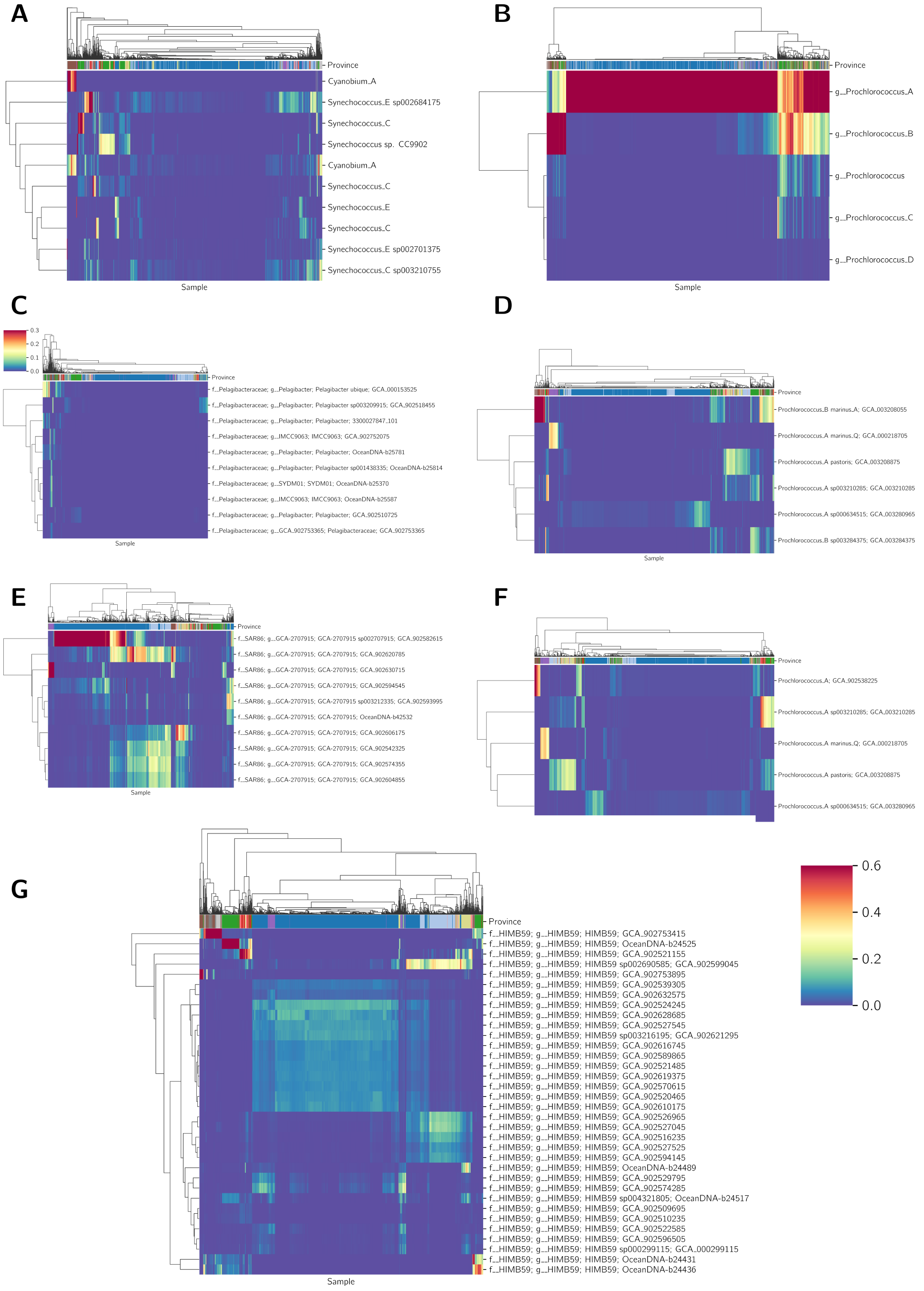


**Supplementary Figure 8. Taxon-specific distributions indicate biogeographic differences in genomic diversity.** Heat maps of relative abundance of most abundant genomes of taxa of interest (rows) across samples (columns). Rows and columns are ordered according to hierarchical clustering with Euclidean distance and average/UPGMA linkage. Dendrograms of hierarchical clustering are displayed along X and Y axes on the top and left side of each panel. Coloured bars at the top indicate province of each sample. Relative abundance is denoted by colour intensity displayed on colour bar on the bottom right, except for C that has its own colour bar. From left to right, top to bottom: genomes *Synechococcus* and *Cyanobium* (A), different genera (as per GTDB classification) of *Prochlorococcus* (B), genomes of *Pelagibacter* (C), most abundant genomes of *Prochlorococcus* genera (D), most abundant genomes of SAR86 (E), most abundant genomes of *Prochlorococcus_A* (high-light lineage) (F), most abundant genomes of HIMB59 (G).

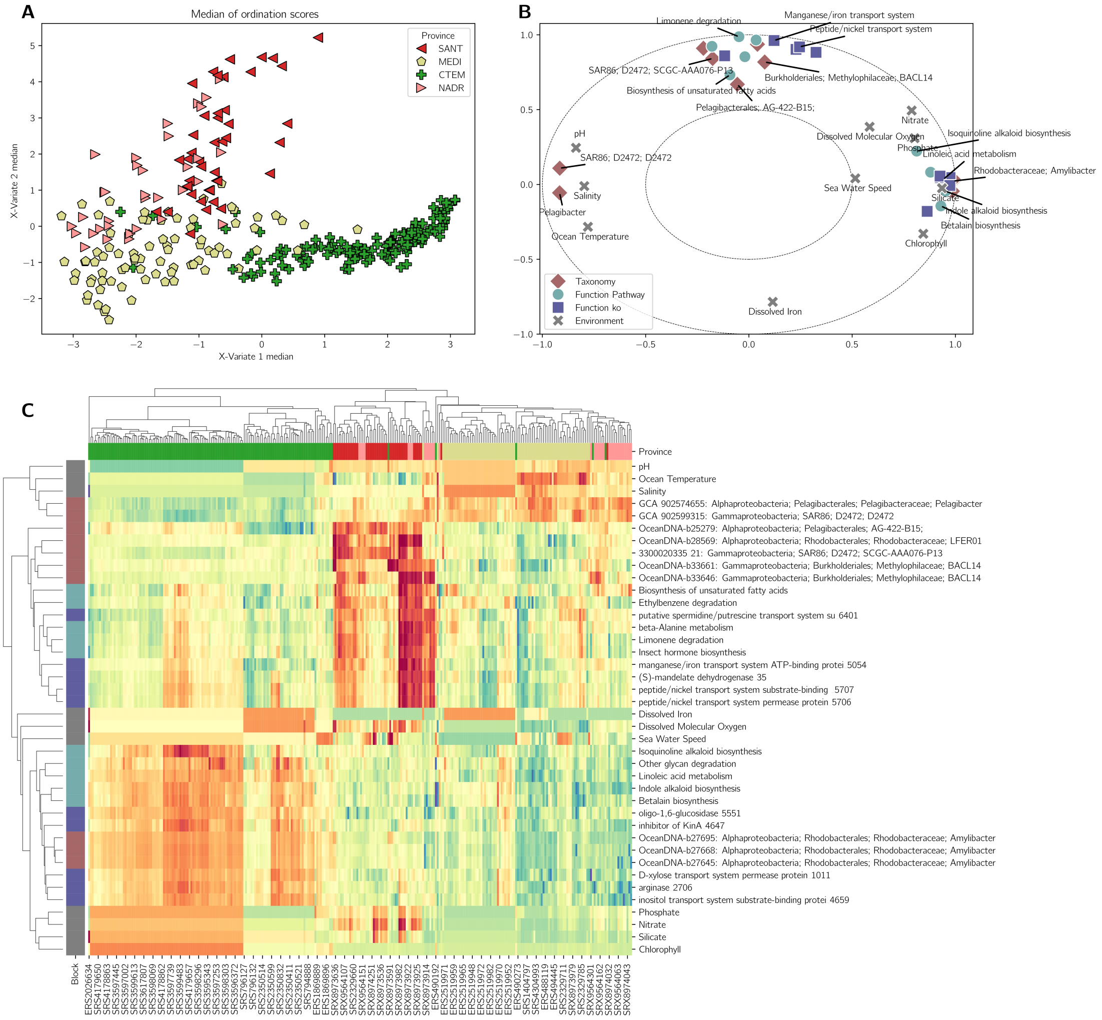


**Supplementary Figure 9. Integrated analysis of temperate provinces.** Block sPLS-DA analysis as described in Figure 5, but for temperate provinces.

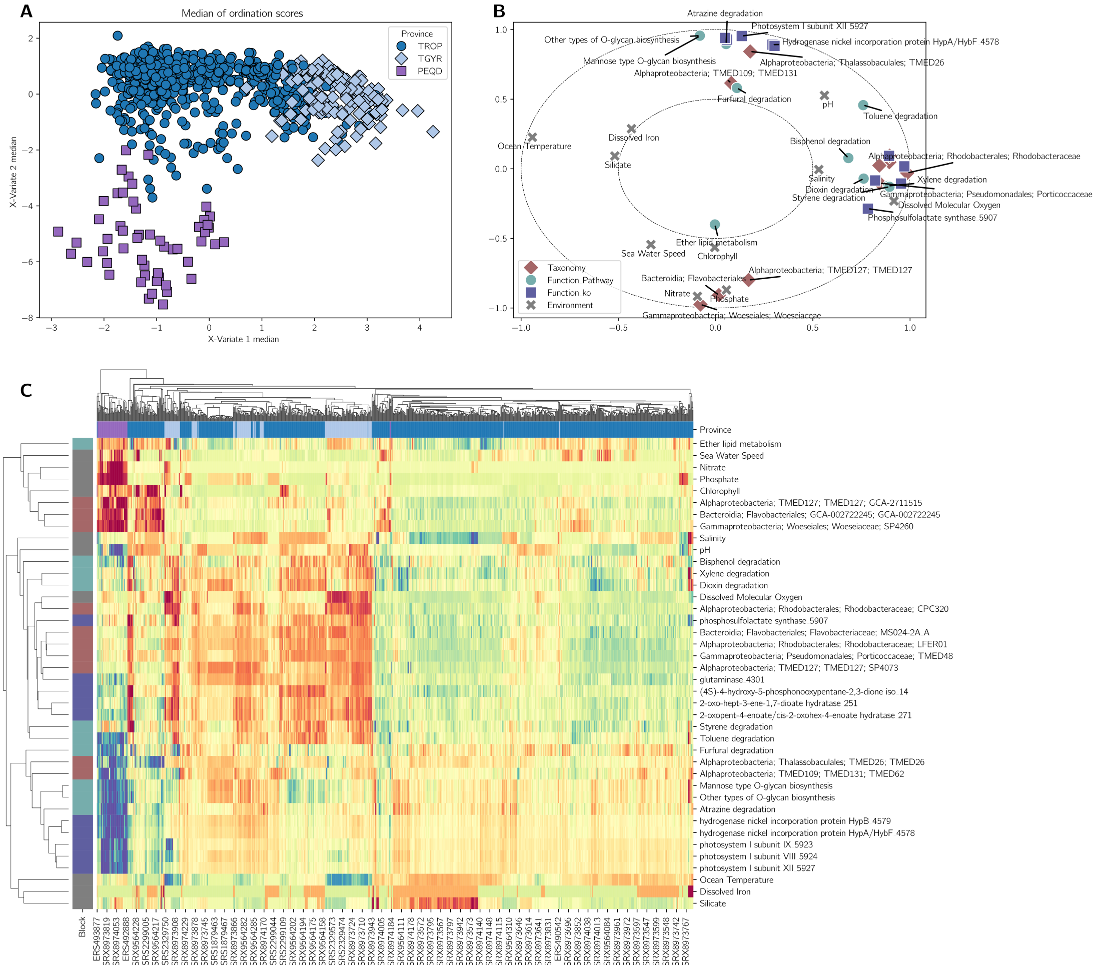

**Supplementary Figure 10. Integrated analysis of tropical provinces.** Block sPLS-DA analysis as described in Figure 5, but for tropical provinces.
